# Silver fluoride as a treatment for the disease dental caries

**DOI:** 10.1101/152207

**Authors:** Jeremy A Horst, Jong Seto

## Abstract

The current paradigm of treatment for dental caries (tooth decay) in primary teeth is dangerous, fails to reach many children, and suffers high recurrence. Acceptance of the paradigm arises from a misperception that untreated caries in primary teeth is a threat to life. We show a linear relationship between age and deaths in the United States from 1999 through 2015 caused by dental caries, pulpal / periapical abscess, or facial cellulitis. The intercept of 6 years coincides with emergence of the first permanent tooth: it appears that caries in primary teeth is not a threat to life. Thus, treatment goals should be to avoid pain, which is not possible with operative dentistry, as it causes pain. Medical management of caries is a distinct treatment philosophy which employs topical minimally invasive therapies that treat the disease, and is not merely prevention. Silver diamine fluoride (SDF) is a central agent to enable effective non-invasive treatment. The announcement of FDA Breakthrough Therapy designation suggests that SDF will become the first FDA approved drug for treating the disease dental caries. Since our last review, 4 clinical trials have been completed, which inform an update to the application protocol and frequency regimen. Suggestions from these studies are to skip the rinsing step due to demonstration of safety and concern of diminished effectiveness by dilution, and to start patients with an intensive regimen of multiple applications over the first few weeks. Breakthroughs in elucidating the impact of SDF on tooth structure and the plaque microbiome inform potential opportunities for bioengineering and understanding caries arrest, respectively. Dentists have been surprised by preference of this treatment over traditional invasive approaches. Renewed interest in this old material has delivered progress to optimize the judicious use of SDF, and enable a revolution in caries management – particularly for primary teeth.

**ONE SENTENCE SUMMARY:** Anesthesia is inappropriate for first-line treatment of early childhood caries now that safe topical treatments such as silver diamine fluoride are available.

## TEXT

### Caries prevalence is nearly complete

Dental caries occurs when dental plaque bacteria ferment dietary sugars into acids that dissolve the tooth. Dental caries is the most prevalent human disease (Murray et al. 2012). >90% of adults are affected by caries (Dye et al. 2015). The modern paradigm of dentistry has reduced disease complications for many. It has not done so for those who have means to access massive amounts of sugar but are sociologically impeded from reducing intake. Modern dentistry treats signs and symptoms of disease, but does not offer a cure (besides full mouth extractions). Further, restorations eventually fail, so nearly always operative treatment progresses from cavitated caries lesion (cavity), to larger filling, to larger yet filling, to crown, to root canal and new crown, to extraction. There are no FDA approved drugs for treating dental caries. Treatment of the disease itself is needed: change the bacteria, strengthen the tooth, enhance the saliva, and change the diet. Medical models of caries treatment attempt to accomplished these goals with antimicrobials, remineralizing agents, salivary stimulation, and behavioral modification.

### Should deadly measures be taken to treat a non-deadly disease?

Young children are increasingly being sedated and anesthetized to enable operative treatment (e.g. fillings; (Bruen et al. 2016), yet there is little evidence that this course of treatment actually makes a significant impact on the disease itself (Twetman and Dhar 2015). The connection between caries in primary teeth and in permanent teeth is weaker than we assume (Heller et al. 2000; Y. Li and Wang 2002). Dental treatment itself inherently causes pain and suffering, which may be greater than that caused by caries. Further, the ubiquitous claim by dentists that untreated dental caries in primary teeth is a threat to life, does not appear to be based on any actual observations.

To assess the potential risk to life by caries in primary teeth, we searched the Centers for Disease Control Wonder database for all causes of any death occurring in the United States (US) during 1999-2015 for ICD-10 codes corresponding to dental caries (K02), pulp / periapical abscess (K04), or cellulitis (K12.2), and stratified by age (Figure 1). The intercept of the relationship between age and death related to caries (6 years of age) coincides perfectly to the average age of eruption of the first permanent tooth. The one death during the age of primary dentition likely corresponds to a 4 year old in New England, known to us, who died from an odontogenic infection of a tooth that was treated under general anesthesia with a pulpotomy and crown. The long-established dental access problem refutes the concept that all children are getting treatment; children with the greatest need for dental care are the least likely to get it. Considering the breadth of these data, and the likelihood of coroners working hard to find actual causes of death in children, this is considerable evidence against a risk of death from untreated dental caries in primary teeth. Meanwhile, the total incidence of death from caries here (58 per year) is similar to dying from lightning strike (51 per year).

**Figure 1.**
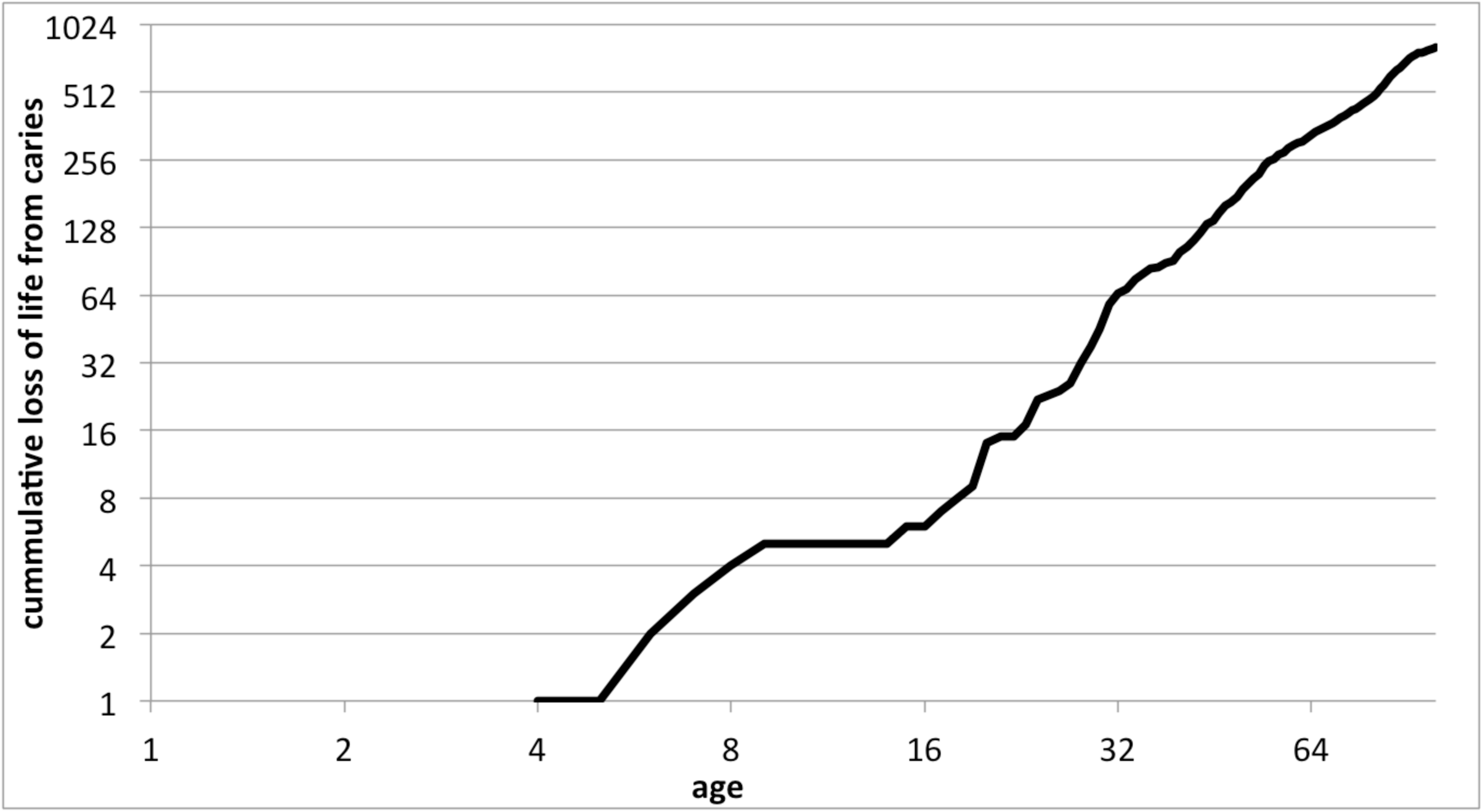
Age versus loss of life caused by dental caries in US. All deaths that occurred in the United States (US) during 1999-2015 with a coroner-assigned cause of death being dental caries (ICD-10 code: K02), pulp / periapical abscess (K04), or cellulitis (K12.2) are plotted by age. The intercept coincides with eruption of the first permanent tooth, suggesting that deaths do not occur from caries in primary teeth.

Irreversible pulpitis in primary teeth can be very painful, and draining abscesses can itch and irritate during eating. Facial cellulitis is obviously painful and concerning, and is treatable with antibiotics. A profound question stems from these data: if caries in primary teeth does not endanger life, is it appropriate to endanger life to treat it? *The treatment must be better than the disease left untreated*. In this context, different treatment goals are indicated: prevent pain and decrease the incidence of new lesions that could cause pain.

### Treatment to achieve prevention

The risk of relapse (recurrent signs of disease; new cavities) following treatment of cavities under general anesthesia (GA) increases from 38±1% (mean ± standard deviation) at 6 months (Primosch et al. 2001; Chase et al. 2004; Berkowitz et al. 2011), to 45±32% at 1 year (Zhan et al. 2006; Hughes et al. 2012), to 62±15% at 2 years (Figure 2; Almeida et al. 2000; Foster et al. 2006; Amin et al. 2010). Patterns of disease without invasive treatment has to our knowledge not been measured. We infer from these data a very low confidence estimate of 38% prevention of any new lesions by traditional operative treatment under GA after 2 years, at most.

**Figure 2.**
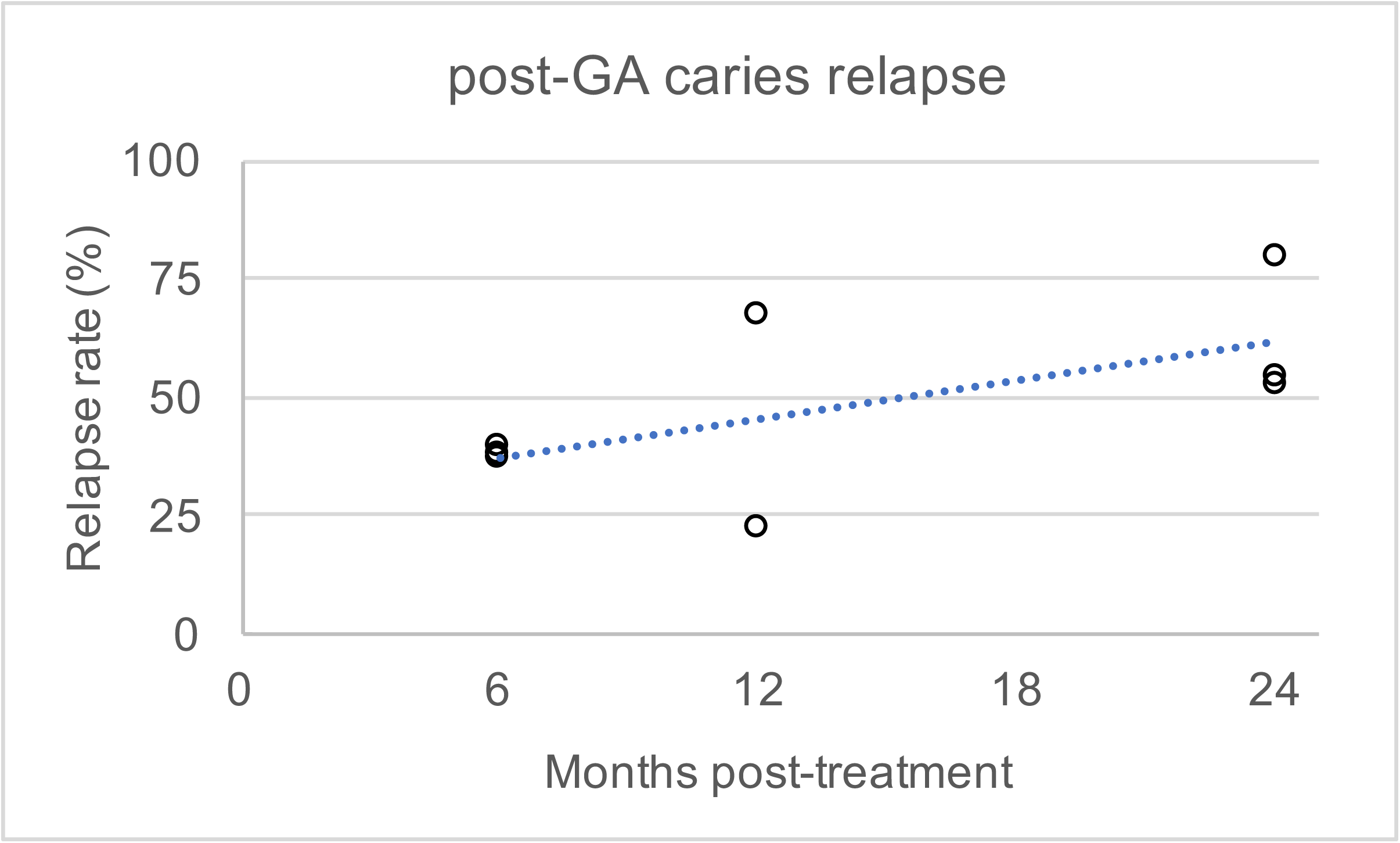
Relapse of signs of dental caries following treatment under general anesthesia. Incidence of new caries lesions following treatment under general anesthesia are plotted against time of evaluation. Linear regression follows y = 1.3x + 29.6, with a correlation coefficient R^2^ = 0.4. Adapted from Twetman and Dhar, 2015, Table 4. References: Almeida et al., 2000; Primosch et al., 2001; Chase et al., 2004; Foster et al., 2006; Zhan et al., 2006; Amin et al., 2010; Berkowitz et al., 2011; Hughes et al., 2012.

Silver diamine fluoride (SDF) is a brush-on liquid that stops 81% of dental caries lesions (Gao, Zhao, et al. 2016). The success rate is similar for restorations placed under GA (Bücher et al. 2014). Stopping lesion progress (caries arrest) appears to have the same effect on preventing pain from the lesion as restoration, but this needs to be studied further. One of the most exciting aspects of SDF is the 67±4% decrease in new lesions on untreated surfaces after 2.5-3 years, achieved simply by treating active lesions (Chu et al. 2002; Llodra et al. 2005). This is not the same as incidence of any new lesions (elaborated above for treatment under GA), however the relation warrants further investigation. The effective treatment of caries lesion sensitivity (Castillo et al. 2011) further indicates SDF as an appropriate treatment for caries. SDF meets the goals of preventing pain and decreasing incidence of new lesions.

### Stopping caries lesion progression (caries arrest)

3 clinical trials on caries arrest by SDF have been published since our systematic review (Figure 3; Horst et al. 2016). One trial in 3-4 year old children documented a dose-response in both application frequency and concentration (Fung et al. 2016). Twice annual application resulted in more arrested lesions after 18 months; similarly 38% SDF (Saforide) stopped more lesions than 12% SDF (Cariostop). The higher effectiveness from increased frequency mimicked that shown previously (Zhi et al. 2012).

**Figure 3.**
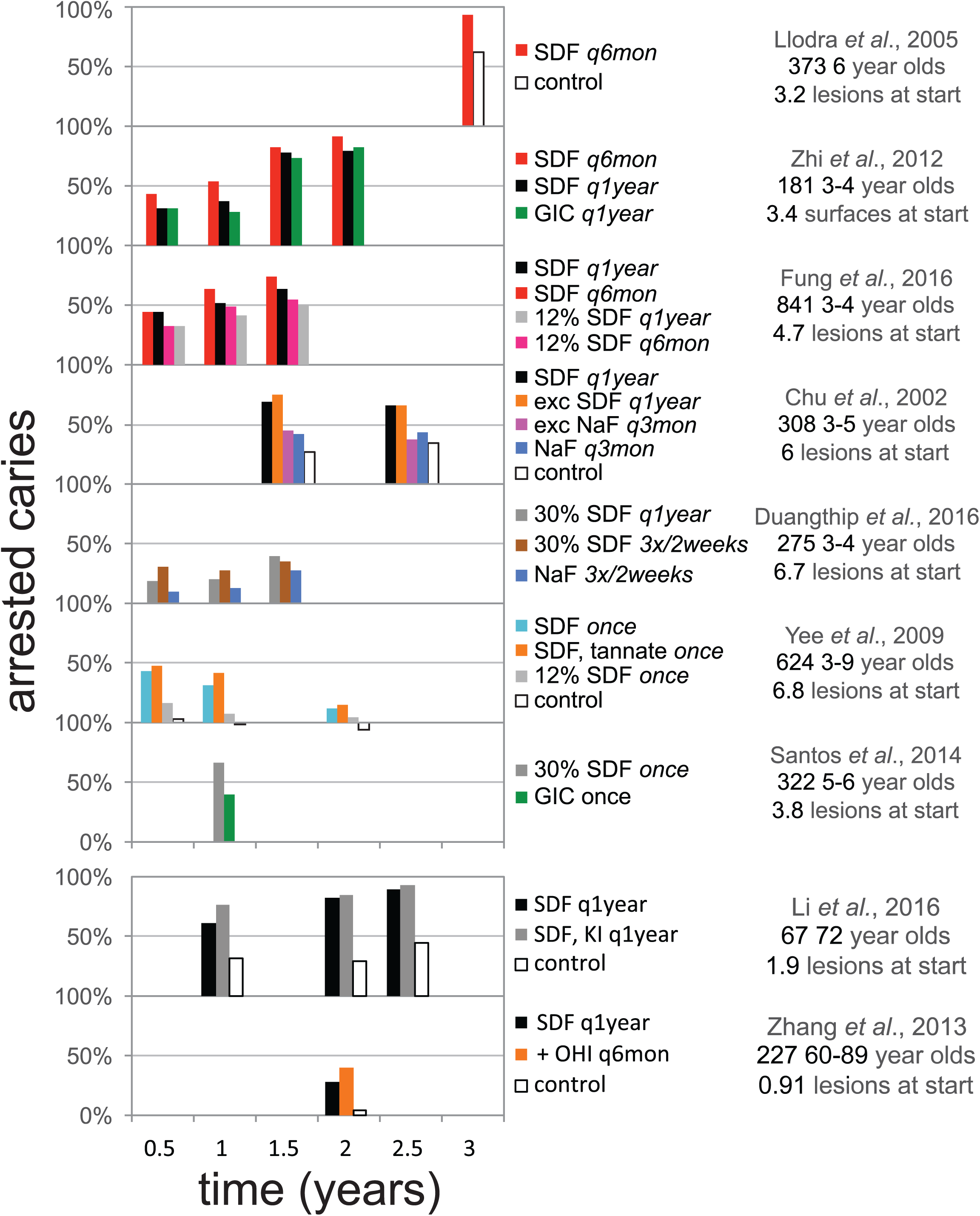
Graphic summary of randomized controlled trials demonstrating caries arrest after topical treatment with SDF. Studies are arranged vertically by frequency of silver diamine fluoride application. Caries arrest is defined as the fraction of initially active carious lesions that became inactive and firm to a dental explorer. SDF (38% unless noted otherwise); q6mon, every six months; q1year, every year; q3mon, every three months; GIC, glass ionomer cement; NaF, 5% sodium fluoride varnish; + OHI q6mon, SDF every year and oral hygiene instructions every six months. Updated from Horst et al., 2016.

Another trial in 3-4 year old children documented increased efficacy at 6 and 12 months following intensive application (3 times in 2 weeks), which was overcome in the single application group by reapplication at 12 months (Duangthip et al. 2016). These outcomes support both the concepts of intensive applications and reapplying over long periods time. It should be noted that much lower arrest rates were seen in this study than others, which may be explained by the concentration of Cariostop actually having ~1/3 of the SDF advertised 30% (Mei et al. 2013).

A trial in adults averaging 72 years of age showed dramatically more effectiveness in arresting caries, 90% (R. Li et al. 2016), than the 28% seen in the one previous study of arrest in older adults (Zhang et al. 2013). This study additionally explored the application of potassium iodide (KI) after SDF, as the interaction of the two produces silver iodide that is yellowish white, instead of black from oxidized silver. This combination did not reduce effectiveness, on the contrary there was a non-statistically significant trend for higher effectiveness at all timepoints. Neither did it reduce staining; the intention of this combination is to decrease color changes from silver oxidation, when remaining sealed and away from light as under opaque glass ionomers (personal communication, Graham Craig). It may be instructive to note that a similar trend in higher effectiveness at all timepoints was also observed following precipitation with tannic acid (Yee et al. 2009).

1,816 patients have been treated with SDF across 12 randomized clinical trials published in English. No significant harms have been noted. This would seem to indicate safety, but in reality, no prospective explicit measure of safety has been published. Too much is at stake to assume. We recently completed a double-blind randomized placebo-controlled superiority trial of SDF in 66 3-5 year old children. We included a safety questionnaire to parents within 48 hours of treatment, and physical assessment at follow up (Milgrom et al. 2017). This “Stopping Cavities” trial documented no adverse events within 17 days after application of blue-tinted SDF without a rinse. Higher levels of arrest were observed in this trial (72%), at 2 weeks versus the common earliest trial outcome of 6 months, which further suggests that effect dissipates with time. Concerns have been expressed about losing effectiveness by rinsing SDF away in the UCSF Protocol; the purpose was concern of safety without it (Horst et al. 2016). The lack of adverse events observed in this study leaves no apparent reason to continue rinsing.

From these 4 trials clinicians may additionally consider intensive application regimens (e.g. 3 times in 2 weeks) and then spreading out applications more over time, skipping the rinse, and further reassurance of: a dose response by application frequency, the need for repeated application over time, and a range from 28-90% arrest in treating root caries in older adults.

### Other non-invasive approaches to arrest caries

While some medicaments decrease the incidence of new lesions, almost no non-invasive therapies available in the US have been shown to stop caries lesions in the dentin. Fluoride varnish reverses 2/3 of enamel lesions (Gao, Zhang, et al. 2016), but makes little impact on dentin caries. While no rigorous clinical trials have yet evaluated silver nitrate, use to treat caries in the early 1900’s was common enough to trust that there is some effectiveness (Black 1908). Sealing in caries, where circumferential enamel is accessible, seems to be the only effective alternative (Mertz-Fairhurst et al. 1998).

Progress against other diseases such as HIV and cancer commonly arise through multi-agent therapy. SDF is simply the combination of an antimicrobial (Ag, 25% w/v), a remineralizing agent (F, 5%), and a stabilizing agent that happens to also be an antiseptic (ammonia, 8%). It is fascinating that none of the components of SDF are highly effective in treating dentinal caries lesions on their own.

### Regulatory progress

In 2014 the FDA cleared silver diamine fluoride (SDF) as a medical device for treating tooth sensitivity. In 2016 the FDA awarded Breakthrough Therapy status as a commitment to an application for approval of SDF as a drug to treat severe early childhood caries. Breakthrough Therapy status does not mean approval, rather it is a commitment to evaluate and assist in the application for a life-threatening disease with no available treatment. Nonetheless, this and the consistent response in many previous clinical trials suggests that SDF will be the first FDA drug to treat dental caries. In 2017, Canada approved SDF with an indication of “anti-caries.” The Indian Health Service released a policy supporting the use of silver ion antimicrobials (SDF or the combination of silver nitrate and fluoride varnish) in their clinics (personal communication, Timothy Ricks). The American Dental Association Council on the Advancement of Access and Prevention has written a resolution in support of use of SDF for caries. The American Academy of Pediatric Dentistry has drafted a policy and guideline supporting use to treat caries as well. This wave of support and interest is appropriate given the many large clinical trials that demonstrate effectiveness.

### SDF adoption

Recent conference presentations described studies that document high levels of acceptance of the stains caused by SDF. An elegant study in New York City asked 33 parents to choose between treatment with SDF or white plastic resin fillings, informing them of the considerations to enable these treatments (Tesoriero and Lee 2017). All parents of “uncooperative” children chose SDF, while 2/3 of parents of other children also chose SDF. A gender disparity emerged, wherein 86% of parents chose SDF for their sons, while only 61% chose SDF for their daughters; still the majority prefer a black stain and uncertainty about outcome over an injection, drill, and prolonged treatment time. The implication is that parents would rather their children have blemishes than experience pain.

Another similar study nearby asked about hypothetical acceptability of the stain. While only 32% of parents accepted SDF for treating anterior teeth initially, a potential requirement of GA to enable operative treatment drove acceptance up to 70% (Crystal et al. 2017). It is interesting to consider how responses might have differed if the studies were conducted after the December 2016 FDA Black Box Warning on the use of GA in pregnancy and before the 3^rd^ birthday. Meanwhile, the vast majority of pediatric dentistry residencies (Nelson et al. 2016), and half of dental school programs are teaching the use of SDF (Ngoc et al. 2017).

### SDF microbial mechanisms

While considerable *in vitro* experiments have documented that SDF inactivates every tested protein and bacterium, no clinical microbiology has been published. The question arises: if SDF kills all bacteria, which microbes are present in the nutrient-rich environment of the SDF-treated caries lesion? To address this question, we performed massively parallel RNA sequencing of a pilot set of plaque samples in the Stopping Cavities trial, taken from 2 caries lesions before and 2 weeks after placebo or control treatments for each child (Milgrom et al. 2017). RNA was used as a proxy for vitality, to enable measurement of all vital microbes; RNA degrades within an hour of production in these conditions. Care was taken to minimize inflow of saliva. The hypothesis was that the relative abundance of caries-associated bacteria would be reduced in the treatment group, but surprisingly, no such changes were observed. Mild increases were seen for only a few bacteria not related to caries, and which pose no known threat. A trend toward increased diversity was seen, rather than the expected decrease that is ubiquitously observed following a course of systemic antibiotics. This signals safety. Abundant high quality RNA was retrieved, which was also surprising. The RNA sequences were additionally scoured for antibiotic or antimetal resistance genes, and these were not changed by treatment. While this was a pilot study in a subset of patients, it is shocking that the microbial composition of the dental plaque on the surface of treated lesions did not significantly change. Meanwhile, our pilot cohort study of 16S microbiome differences found differential abundance of *Streptococcus mutans* when lesions failed to arrest after SDF treatment (*unpublished data*).

### SDF structural mechanisms

The outer surface of SDF-arrested caries lesions is approximately twice as hard as normal dentin (Chu and Lo 2008). The assumption that SDF achieves this by fluoride-mediated remineralization is not reasonable; the collagenous matrix of the outermost dentin in a carious lesion is far too degraded to be remineralized. Yet no structural observations have explained the hardness of SDF treated lesions. To address this disparity, we evaluated clinically treated, exfoliated or extracted teeth using synchrotron microCT. The advantage of the synchrotron source is essentially the abundance of energy, such that we can filter out X-rays that do not help differentiate between densities in the range of tooth structures and solid silver, and be left with plenty of X-rays with which to make images. These experiments revealed solid wires microns in diameter, coursing millimeters through dentinal tubules. We verified the composition of these wires as silver using SEM-EDX, and documented the consistent presence of these “microwires” across every imaged tooth (*unpublished data*). An example of these wires in a permanent tooth treated clinically is shown in Figure 4. The silver microwires likely provide the same physical reinforcement to weakened dentin as rebar does in cement, ending the mystery of lesion hardening.

**Figure 4.**
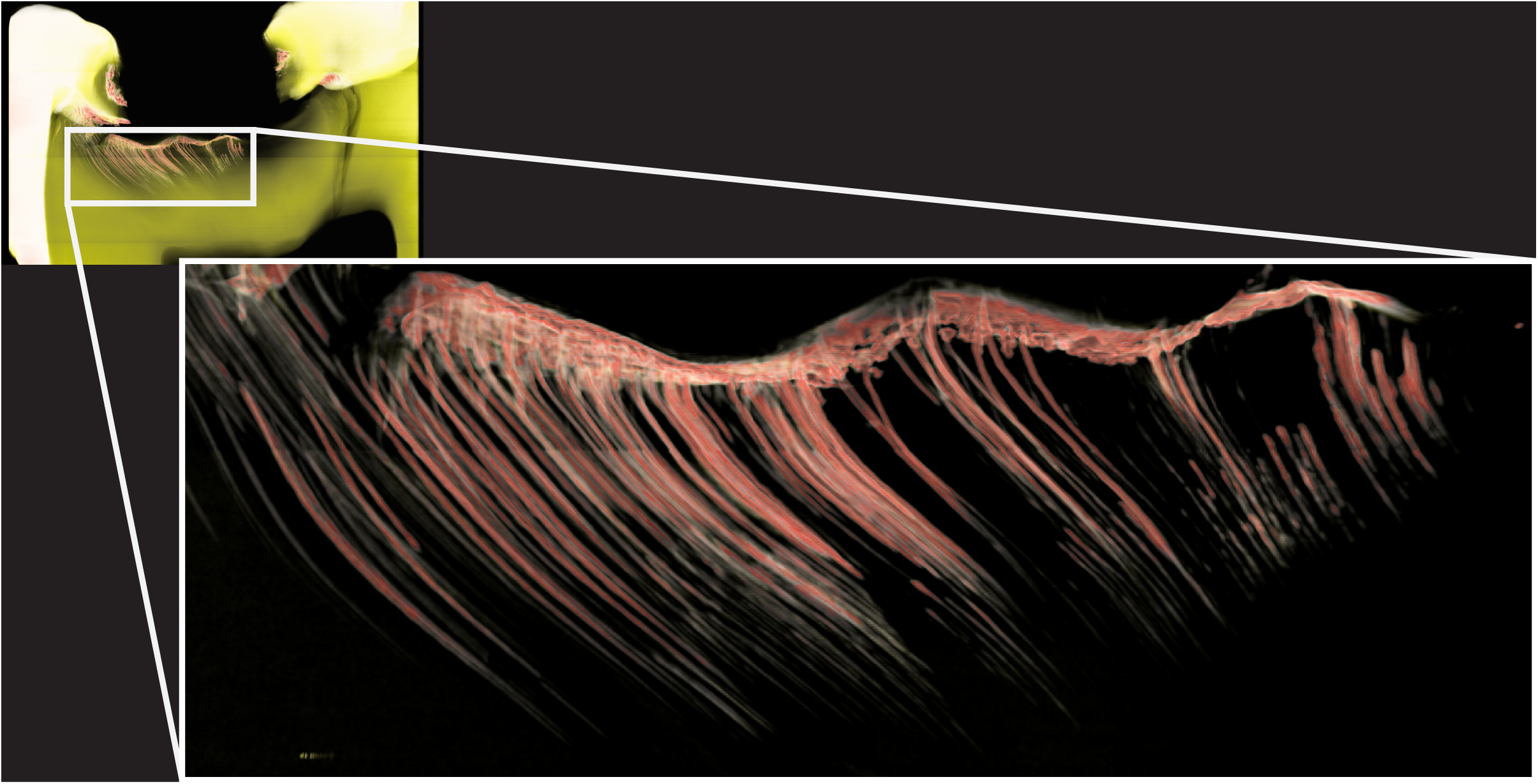
Silver microwires coursing through dentinal tubules following clinical treatment of caries lesions with SDF. Image produced by synchrotron microCT. White corresponds to the density range of enamel, yellow to that of dentin, red denoting density significantly above that of enamel, and black being the highest density observed.

### Summary

The appropriateness of traditional operative dentistry as the first line of treatment for dental caries is in question. We show evidence for absence of any fatal risk from caries in primary teeth. This, and the FDA Black Box Warning against general anesthetics in young children, urge a paradigm shift. Clearance of SDF in the United States provides an agent for change to non-invasive caries management. Rapid adoption despite the non-esthetic results indicates preference against the discomfort required by traditional operative dentistry, which is supported by cursory surveys. New clinical trial data suggests starting with more frequent applications and decreasing frequency with time, while maintaining at least annual application, and removing the rinse step. Our recent work documents a surprising lack of changes to the dental plaque microbiota following SDF treatment, and that SDF produces solid silver microwires, cast *in situ*, which explain the increased lesion hardness and may inform mechanistic treatment goals. While more work needs to be done to understand and anticipate treatment failure, our mechanistic understanding of caries arrest is starting to catch up with the massive clinical trial data – all of which point to effectiveness and safety for treatment of dental caries by SDF.

## ACKNOWLEDGEMENT

This work was supported by NIH NIDCR grant T32-DE007306. Thanks to Jason Hirsch, John Frachella, Steve Duffin, Jeanette MacLean, and Peter Milgrom for thoughtful discussions on the clinical use of SDF for treating dental caries. The authors declare that no conflicting financial, personal, or professional interests have influenced this work.

